# Cytoduction preserves genetic diversity following plasmid transfer into pooled yeast libraries

**DOI:** 10.1101/2024.05.24.595802

**Authors:** Han-Ying Jhuang, Dimitra Aggeli, Gregory I. Lang

## Abstract

Much of our understanding of functional genomics derives from insights gained from large strain libraries including the yeast deletion collection, the GFP and TAP-tagged libraries, QTL mapping populations, among others [1-5]. A limitation of these libraries is that it is not easy to introduce reporters or make genetic perturbations to all strains in these collections. Tools such as Synthetic Genetic Arrays allow for the genetic manipulation of these libraries but are labor intensive and require specialized equipment for high throughput pinning [6]. Manipulating a diverse library *en mass* without losing diversity remains challenging. Ultimately, this limitation stems from the inefficiency of transformation, which is the standard method for genetic manipulation in yeast. Here, we develop a method that uses cytoduction (mating without nuclear fusion) to transfer plasmids directionally from a “Donor” to a diverse pool of “Recipient” strains. Because cytoduction uses mating, it is a natural process and is orders-of-magnitude more efficient than transformation, enabling the introduction of plasmids into high-diversity libraries with minimal impact on the diversity of the population.

## Results & Discussion

The mating process in *Saccharomyces cerevisiae* involves the fusion of haploid yeast cells of opposite mating types (*MAT***a** and *MAT*_α_) to form a *MAT***a/**_α_ diploid. If one of the mating partners carries the *kar1*Δ*15* mutation, mating will result in cell fusion but not nuclear fusion [7]. This process (cytoduction) is an efficient method for transferring plasmids, mitochondria, cytoplasmic viruses, prions, and rarely, individual chromosomes [8, 9]. To transfer a plasmid to a diverse library of Recipient strains, we first introduce the plasmid into a Donor (*kar1*Δ*15* mutant) strain. We then mate the Donor strain to the Recipient library. Following mating, we select for the Recipient library and the transferred plasmid, and we counter-select against the Donor strain and Donor/Recipient diploids.

In this study, we set up a direct comparison between transformation and cytoduction as a means of introducing plasmids into yeast libraries. For this test we used a barcoded library of segregants from a cross between BY4742 and RM-11a [10]. This library contains 4,401 genotypes, each carrying a unique barcode. We conducted parallel cytoduction and transformation experiments using four different plasmids. These plasmids were either CEN/ARS-based or 2-micron-based and included a selectable marker, either KanMX or NatMX. The plasmids were introduced to the Recipient library through either transformation or cytoduction. We conducted the transformation and cytoduction experiments using an equal division (∼10^7^ cells each) of the same initial population. Transformations were performed using the standard lithium acetate protocol with 1 μg of plasmid. Cytoductions were performed by mixing the *MAT***a** Recipient library with a five-fold excess of *MAT*_α_ Donor cells, followed by six hours of mating on YPD before selective plating. We collected the selected cells (∼10^5^ colonies for each of the transformations and ∼10^7^ cells for each of the cytoduction experiments, at least two orders of magnitude above the library diversity) and amplified barcodes for Illumina sequencing, using ∼10^8^ cells per experiment, which were then sequenced to a depth of ∼5x10^5^ reads per sample.

Following cytoduction, most barcodes from the initial pool were recovered (97% on average), and these exhibited a strong correlation with the initial pool (0.88 on average, Figure 1A and Supplemental Figure 1A). In contrast, after transformation, only 43% of the barcodes were recovered in the final pools on average, and these exhibited a lower correlation with the initial pool (0.49 on average). As a result of preserved barcode frequencies, fewer sequencing reads are needed to recover a greater number of barcodes from cytoduction compared to transformation experiments (Figure 1B).

**Figure 1.**
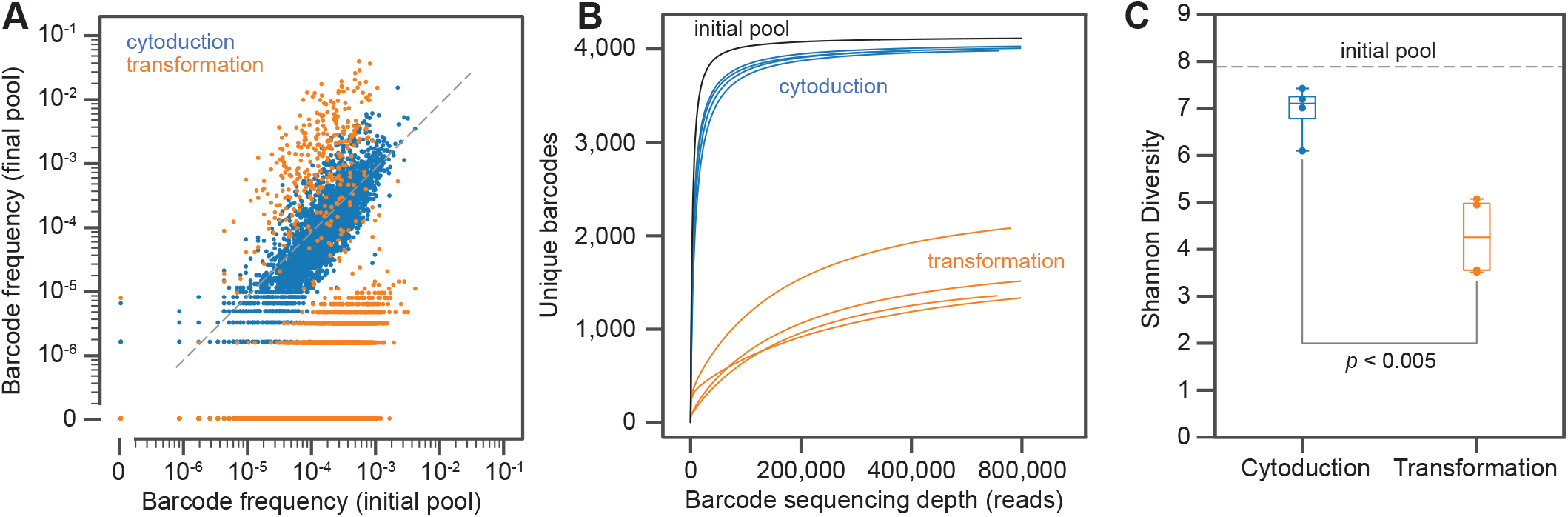
Cytoduction is highly efficient at transferring plasmids while maintaining diversity in the Recipient population. We introduced four plasmids representing all combinations of 2-micron or CEN/ARS plasmids with NatMX or KanMX markers into ∼10^7^ cells from a library containing 4,401 unique barcodes. A total of 4,149 barcodes were identified in the input library. **(A)** The distribution of barcode frequencies shows that cytoduction maintains relative abundance. The data shown are from the cytoduction and transformation experiments using the KanMX-marked CEN/ARS plasmid. The barcode recovery and correlation coefficients are 98%/0.88 and 38%/0.41 for cytoduction and transformation, respectively. The other three scatter plots are shown in Supplemental Figure 1A. **(B)** Rarefaction curves show that cytoduction recovers significantly more barcodes compared to transformation, especially with fewer than 100k reads. Even with >400k reads, ∼60% of the barcodes are missing from the transformation, but only ∼2.5% from the cytoduction. **(C)** Shannon Diversity is higher in the cytoduction population compared to transformation across all four experiments (*p* < 0.005, paired t-test).

We quantified barcode diversity using Shannon entropy, a standard metric used to assess diversity. Shannon entropy was the highest for the input pool with only a minor loss of diversity following cytoduction (Figure 1C). In contrast, we observed a substantial loss of diversity following transformation (Figure 1C, *p* < 0.005, paired t-test), indicating that transformation is comparatively low efficiency. We do not find significant differences in Shannon entropy when comparing experiments with different plasmid types, indicating that efficiency is not strongly influenced by the selective marker or the type of replication origin on the plasmid (Supplemental Figure 1B, *p* = 0.041 and *p* = 0.275, paired t-tests).

To test the extent to which genetic factors influence the observed changes in barcode frequencies, we performed a QTL analysis using the known genotypes associated with each barcode. Fitness values were estimated based on changes in barcode frequencies as described previously [10]. We then calculated the correlation between fitness and the genotype at each of the 41,595 polymorphic sites (Supplemental Figure 2). For cytoduction, we observed enrichment for loci commonly associated with growth but not mating efficiency. This suggests that the five-fold excess of Donor strain mitigated selection on mating but that the six-hour outgrowth may impose modest selection for loci effecting growth rate. For transformation, in contrast, we observed few enriched loci, suggesting that sampling is largely stochastic and not strongly influenced by genetic variation represented in our library.

This study demonstrates the efficacy of cytoduction as a tool to introduce plasmids into yeast libraries while preserving the diversity and the relative abundance of individual genotypes. We were able to manipulate 97% of the variants, while also achieving good population representation (correlation with the initial pool is 0.88). As a comparison, only 43% of the variants were retained after chemical transformation with correlation being only 0.49. At a minimum the method requires a *kar1*Δ*15* Donor strain and appropriate genetic markers to select for the haploid Recipient library and/or against diploid and Donor cells. This method could be combined with CRISPR/Cas9 editing, further expanding the potential for modifying the genetic background of diverse yeast libraries.

## Acknowledgements

We thank members of the Lang Lab for comments on the manuscript. This study was supported by a grant from the National Institutes of Health (R35GM149540). Portions of this research were conducted on Lehigh University’s Research Computing infrastructure partially supported by the National Science Foundation (Award 2019035).

---

**Supplemental Figure 1.**
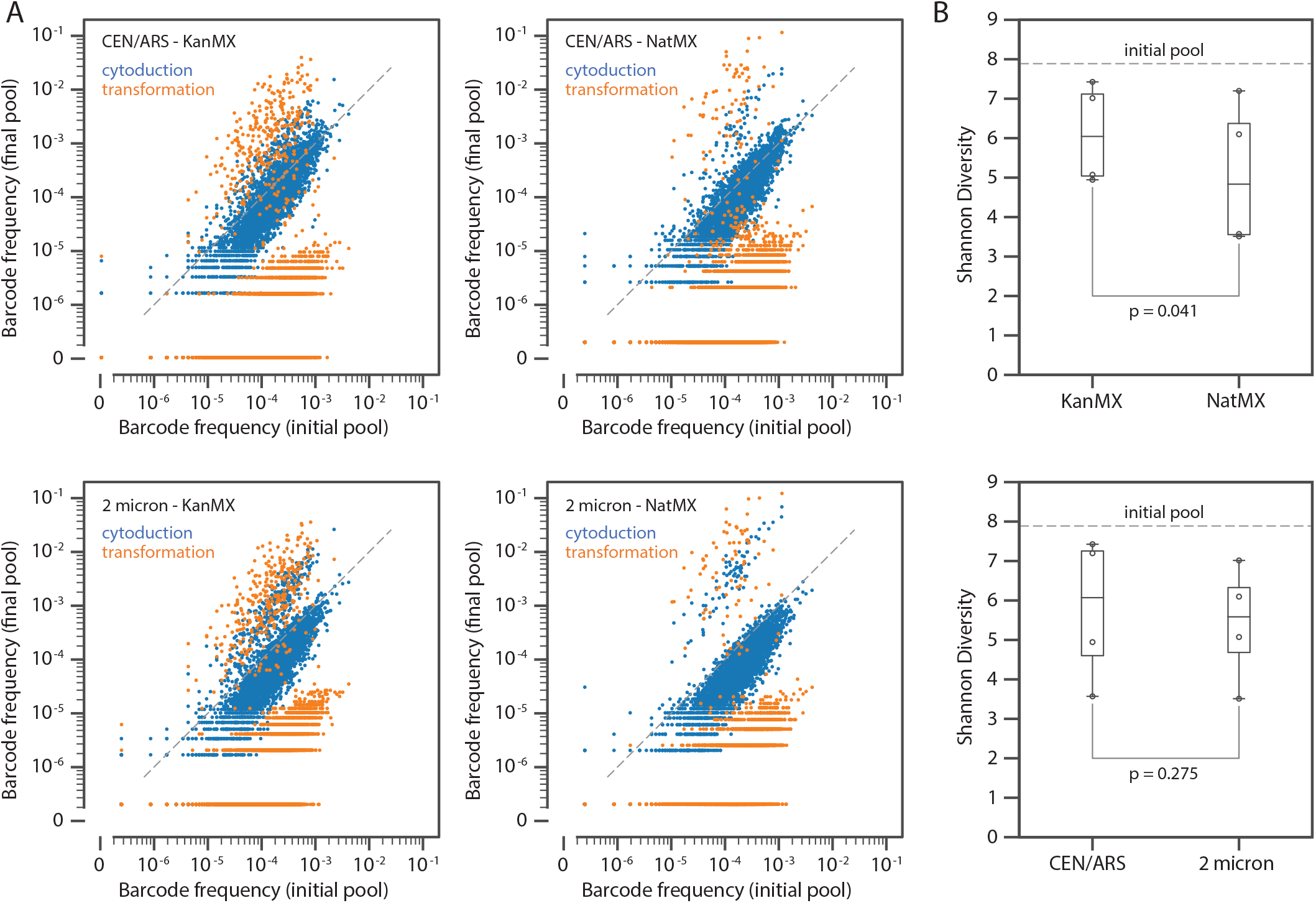

**Supplemental Figure 2.**
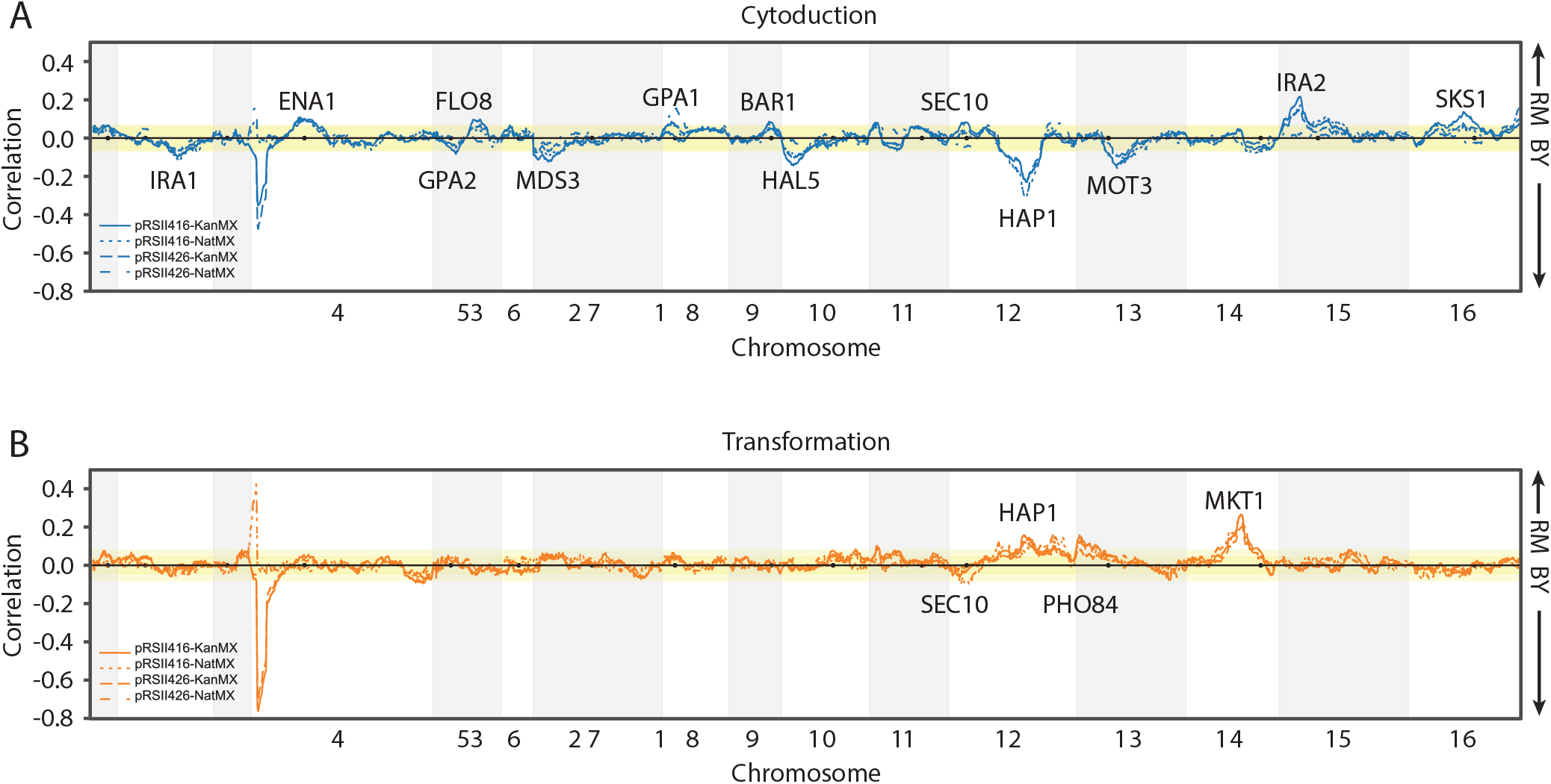

