## Supplemental Methods and Figure Legends for "Cytoduction preserves genetic diversity following plasmid transfer into pooled yeast libraries"

**Materials and Methods**

### *Donor and Recipient Strains*

We used a pooled library of 4,401 barcoded segregants from a cross between BY4742 and RM11-1a strains [1]. The library has the genotype *MAT***a** *ura3∆ can1∆::pSTE2-SpHIS5 ho∆::URA3* and the barcodes are within an artificial intron in *URA3*. We propagated the population and archived multiple aliquots at cell density 10^8^ cells/ml. A single aliquot was thawed and 200 μl was used to inoculate 10 ml YPD (Yeast Extract, Peptone, Dextrose), which was grown for 10 hours at 30^o^C. The culture was then diluted 1:20 in YPD, incubated for 3 hours at 30^o^C and the resulting log-phase cells were used for the transformation and cytoduction experiments described below. Throughout the growth, selection, and genomic and library preparations, population bottlenecks were at least eight-fold larger than the library diversity. The starting population lineage frequencies spanned four orders of magnitude (Supplemental Figure 1A).

The Donor strains for cytoduction were built upon yGIL1940, an S288c derivative with the genotype *MAT*α *Hap1 leu2∆ his3∆ met17∆ nej1∆ ura3∆ CAN1 kar1∆15*. yGIL1940 was transformed with four plasmids: pRSII416-KanMX, pRSII416-NatMX, pRSII426-KanMX, and pRSII426-NatMX. All four plasmids carry the *URA3* marker. These plasmids have the four possible combinations of drug marker (KanMX and NatMX) and base (2-micron and CEN/ARS).

### *Chemical Transformation*

The Recipient yeast population was transformed using the lithium acetate protocol [2] with either of four plasmids – pRSII416-KanMX, pRSII416-NatMX, pRSII426-KanMX, and pRSII426-NatMX. For the transformation we used ~10^7^ mid-log phase Recipient cells and 1 µg of plasmid DNA. Heat shock at 42˚C was done for 25 min. Before selection on 30-cm agar plates, cells were recovered in YPD at 30˚C for an hour to allow for drug resistance expression. SD-HIS+G418 plates were used for pRSII416-KanMX and pRSII426-KanMX, and SD-HIS+ClonNAT plates were used for pRSII416-NatMX and pRSII426-NatMX, respectively. Approximately 10^5^ colonies were harvested by scraping after incubation at 30˚C for 2 days and subjected to genomic DNA extraction.

### *Plasmid Transfer by Cytoduction*

We performed cytoduction by mixing ~10^7^ mid-log phase cells of the yeast library with ~5x10^7^ mid-log phase cells of each of the Donor strains in 10 ml final volume of YPD. The mixtures were then concentrated on filter paper using a vacuum manifold. The filters were transferred to YPD plates and incubated at 30˚C for 6 hours to allow for ~1 round of mating. Mating mixtures were then washed off in 1 ml of sterile distilled water and spread on selection plates. The same selective medium was used for both the transformation and cytoduction – SD-HIS+G418 plates were used for pRSII416-KanMX and pRSII426-KanMX, and SD-HIS+ClonNAT plates were used for pRSII416-NatMX and pRSII426-NatMX, respectively. Cells were harvested by scraping after incubation at 30˚C for 2 days and subjected to genomic DNA extraction. The number of colonies far exceeded the number of colonies yielded by transformation, as cytoduction is much more efficient.

### *Barcode Sequencing*

The libraries were prepared as previously described [3]. Genomic DNA extraction was done as described previously using 10^8^ cells per sample and quantified by Qubit. Briefly, 1 µg of DNA was used in a 2-step PCR protocol to amplify the barcoded locus and introduce Unique Molecular Identifiers (UMIs). DNA from all libraries was pooled isostoichiometrically based on DNA concentrations estimated by Nanodrop. A 350-bp band was gel purified with the QIAGEN gel extraction kit (QIAGEN, Germantown, MD). Final sample quantification was done by Qubit. Final pools were analyzed by BioAnalyzer on a High-Sensitivity DNA Chip (BioAnalyzer 2100, Agilent) before Illumina sequencing on a NovaSeq with 150x150 bp paired-end reads at the Sequencing Core Facility within the Lewis-Sigler Institute for Integrative Genomics at Princeton University. Each library had at least 5 x 10^5^ reads (Figure 1B). Barcodes were extracted from raw reads using UMI-tools, which uses network-based methods to account for errors [4]. A list of known barcodes was given, and a pattern that allows up to 3 mismatches in each of the 2 invariable regions around the barcodes was used for regex search of the barcodes. As a result, 92.3±0.8% of the reads matched barcode sequences. Estimates for the required number of reads for recovering barcodes after cytoduction and transformation were obtained by downsampling. We quantified barcode diversity using Shannon entropy, a common metric for assessing population diversity [5].

### *QTL Mapping*

We performed a genome-wide scan for the QTL using Pearson correlation. First, we used inferred probabilistic genotype values and inferred fitness values with a likelihood model for time-dependent barcode frequencies [1]. As the genotype values are probabilities and not definitive, we next calculated the Pearson correlation coefficient between the inferred phenotype and genotype values at each locus using a linear regression model. Standard error of correlation coefficients across the genome and experiments was used to find significant calls. Correlation values greater than one standard error were considered significant. Annotations are candidate genes based on previously reported QTL that colocalize with peaks in the correlation values [1]. The prominent QTL on Chromosome IV are due to the barcode locus [1]. KanMX and NatMX were used to mark the barcode locus in BY and RM, respectively. Though the markers should have been removed during construction, it is likely that some segregants retained the marker. Selections on G418 and ClonNat enriched for segregants carrying BY or RM parentage, respectively, around the barcode locus.

**Supplementary Figure Legends**

**Supplemental Figure 1. Cytoduction and transformation efficiencies do not depend on plasmid type or selectable marker. (A)** The distribution of barcode frequencies shows that cytoduction maintains relative abundance and transformation does not. The upper left panel is the same as in Figure 1A. The other three panels show that this result holds for four plasmids tested. The barcode recovery and correlation coefficients for cytoduction are 98%/0.88, 97%/0.89, 98%/0.85, and 97%/0.88 for CEN/ARS KanMX, CEN/ARS NatMX, 2 micron KanMX, and 2 micron NatMX, respectively. The barcode recovery and correlation coefficients for transformation are 38%/0.41, 41%/0.51, 55%/0.55, and 36%/0.49 for CEN/ARS KanMX, CEN/ARS NatMX, 2 micron KanMX, and 2 micron NatMX, respectively. **(B)** Shannon diversity is not impacted by the plasmid marker (KanMX or NatMX) or plasmid type (CEN/ARS vs 2-micron). Shannon diversity values were compared by a paired t-test.

**Supplemental Figure 2. Changes of lineage frequencies in cytoduction correlate with loci associated with growth while few loci were correlated in transformation. (A)** Correlation between individual polymorphic sites and changes of lineage frequencies in cytoduction. Positions of the peaks colocalize with genes associated with growth in the previous study [1]. **(B)** Correlation between individual polymorphic sites and changes of lineage frequencies in transformation. Fewer QTL are enriched in the transformation experiments, suggesting random sampling as the predominant factor for the changes in barcode frequencies.
